# PERK-mediated antioxidant response is key for pathogen persistence in ticks

**DOI:** 10.1101/2023.05.30.542958

**Authors:** Kristin L. Rosche, Joanna Hurtado, Elis A. Fisk, Kaylee A. Vosbigian, Ashley L. Warren, Lindsay C. Sidak-Loftis, Sarah J. Wright, Elisabeth Ramirez-Zepp, Jason M. Park, Dana K. Shaw

## Abstract

A crucial phase in the lifecycle of tick-borne pathogens is the time spent colonizing and persisting within the arthropod. Tick immunity is emerging as a key force shaping how transmissible pathogens interact with the vector. How pathogens remain in the tick despite immunological pressure remains unknown. In persistently infected *Ixodes scapularis*, we found that *Borrelia burgdorferi* (Lyme disease) and *Anaplasma phagocytophilum* (granulocytic anaplasmosis) activate a cellular stress pathway mediated by the endoplasmic reticulum receptor PERK and the central regulatory molecule, eIF2α. Disabling the PERK pathway through pharmacological inhibition and RNAi significantly decreased microbial numbers. *In vivo* RNA interference of the PERK pathway not only reduced the number of *A. phagocytophilum* and *B. burgdorferi* colonizing larvae after a bloodmeal, but also significantly reduced the number of bacteria that survive the molt. An investigation into PERK pathway-regulated targets revealed that *A. phagocytophilum* and *B. burgdorferi* induce activity of the antioxidant response regulator, Nrf2. Tick cells deficient for *nrf2* expression or PERK signaling showed accumulation of reactive oxygen and nitrogen species in addition to reduced microbial survival. Supplementation with antioxidants rescued the microbicidal phenotype caused by blocking the PERK pathway. Altogether, our study demonstrates that the *Ixodes* PERK pathway is activated by transmissible microbes and facilitates persistence in the arthropod by potentiating an Nrf2-regulated antioxidant environment.

## INTRODUCTION

Ticks are prolific spreaders of pathogens that plague human and animal health including bacteria, viruses, and protozoan parasites^1–4^. A crucial phase in the tick-borne pathogen lifecycle is the time spent colonizing and persisting within the arthropod vector^5^. While many forces impact the way transmissible pathogens interface with their arthropod vectors, recent advances have demonstrated that tick immunity is an important influence shaping this interaction. Immune functions that respond to tick transmitted bacterial pathogens include cellular defenses, such as phagocytosis by hemocytes, and humoral defenses orchestrated by the IMD (Immune Deficiency) and JAK-STAT (Janus kinase-signal transducer and activator of transcription) pathways^6–17^. Notably, the tick IMD pathway is divergent from what has canonically been described in *Drosophila*. Ticks and other non-insect arthropods lack genes encoding key molecules such as transmembrane peptidoglycan recognition proteins that initiate the IMD pathway, and the signaling molecules *IMD* and *FADD*^10, 18, 19^. Instead, the tick IMD pathway responds to multiple cues such as infection-derived lipids that are sensed by the receptor Croquemort^10, 11, 17^ and to cellular stress that is caused by infection^20, 21^.

Recently, the unfolded protein response (UPR) has been linked to arthropod immunity^20^. The UPR is a specialized cellular response pathway that is activated when the endoplasmic reticulum (ER) is under stress^22–24^. Three ER receptors orchestrate the UPR and function to restore cellular homeostasis: ATF6 (activating transcription factor 6), PERK (PKR-like ER kinase), and IRE1α (inositol-requiring enzyme 1α). When *Ixodes scapularis* ticks are colonized by *Borrelia burgdorferi* (Lyme disease) or *Anaplasma phagocytophilum* (granulocytic anaplasmosis), the IRE1α receptor undergoes self-phosphorylation and pairs with TRAF2 (TNF receptor associated factor 2) to activate the IMD pathway^20^. During this process, reactive oxygen species (ROS) are also potentiated. This signaling network functionally restricts the number of *Borrelia* and *Anaplasma* that colonize the tick^20^. Furthermore, the UPR-IMD pathway connection and its pathogen restricting potential is present in several arthropods against multiple types of pathogens, suggesting that this signaling network may be an ancient mode of pathogen-sensing and vector defense against infection^20^.

As vector immunity continues to be explored, a fundamental question has emerged: how are tick-borne pathogens persisting in the arthropod despite immunological pressure? Herein, we report that *B. burgdorferi* and *A. phagocytophilum* trigger phosphorylation of the central regulatory molecule, eIF2α, in *I. scapularis* ticks through the ER stress receptor PERK. Knocking down the PERK-eIF2α-ATF4 pathway *in vivo* through RNAi significantly inhibited *A. phagocytophilum* and *B. burgdorferi* colonization in ticks and reduced the number of microbes persisting through the molt. Infection-induced PERK pathway activation in *Ixodes* was connected to the antioxidant transcription factor, Nrf2. Disabling Nrf2 or the PERK pathway in tick cells caused accumulation of ROS and reactive nitrogen species (RNS) that led to greater microbial killing. This microbicidal phenotype could be rescued by exogenously supplementing antioxidants, demonstrating that the PERK pathway supports microbial persistence by detoxifying ROS/RNS. Overall, we have uncovered a mechanism at the vector-pathogen interface that promotes persistence of transmissible microbes in the arthropod despite active immune assaults.

## RESULTS

### Cellular stress genes are transcriptionally induced in infected, unfed I. scapularis nymphs

Infectious microbes impart cellular stress on the host^25^. For this reason, we investigated whether cellular stress responses impact how microbes survive in ticks^20^. We previously observed that the IRE1α-TRAF2 axis of the *I. scapularis* UPR responds to *A. phagocytophilum* and *B. burgdorferi* and functionally restricts pathogen colonization during a larval blood meal by crosstalking with the IMD pathway and potentiating ROS (Fig 1A)^20^. How *Anaplasma* and *Borrelia* persist in the tick despite this immunological pressure is not well-understood. In this study, we analyzed the transcriptional response of *I. scapularis* nymphs that were infected but were unfed (flat) to explore how ticks respond to persistent infection. We found that, similar to results from immediately repleted ticks^20^, unfed nymphs that are infected with *A. phagocytophilum* or *B. burgdorferi* showed increased expression of genes associated with IRE1α-TRAF2 signaling (Fig 1B-D). In addition, we also found increased expression of genes that are part of the PERK pathway and another cellular stress response network termed the “integrated stress response” (ISR) (Fig 1E-I).

**Figure 1.**
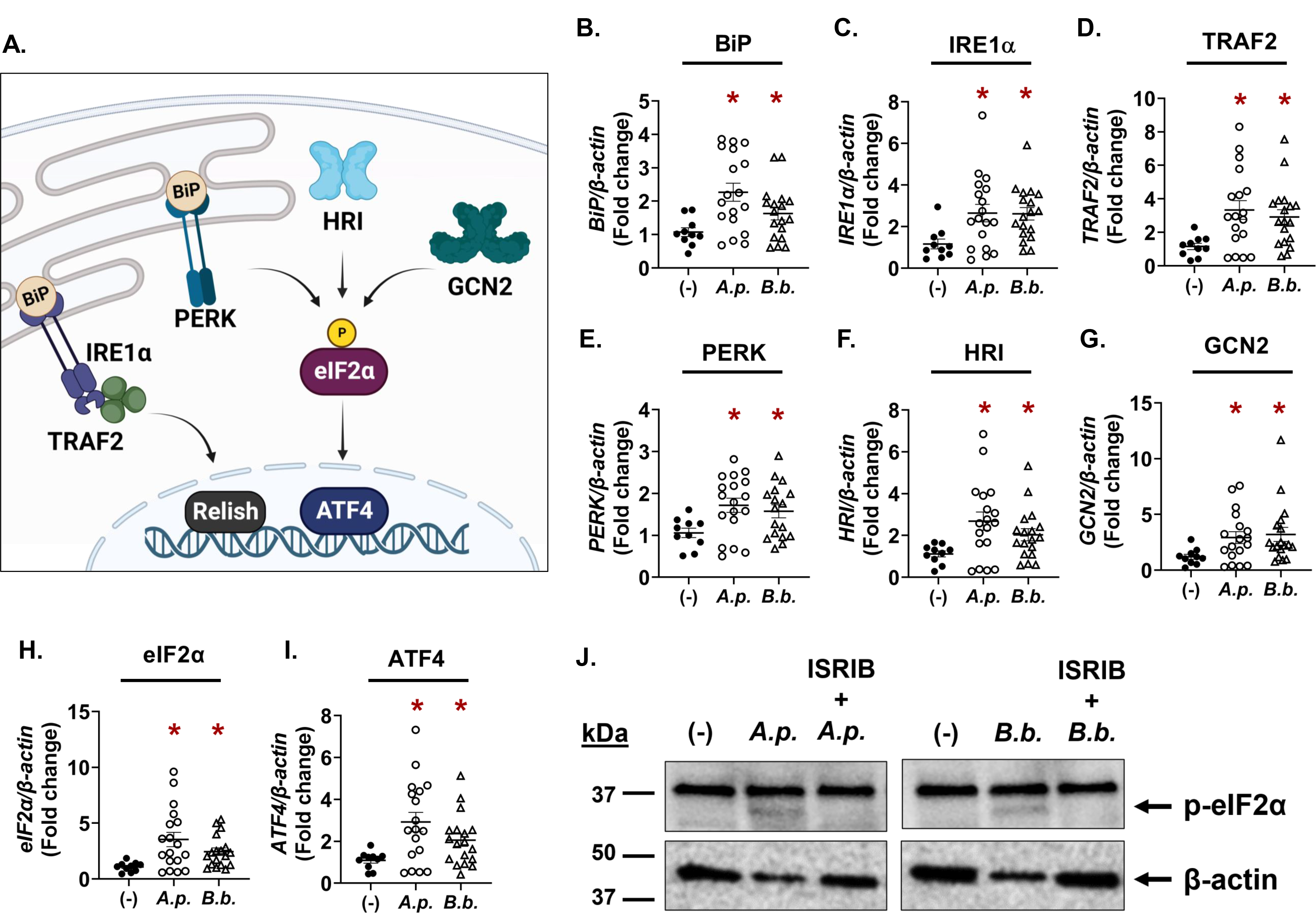
**Tick-borne pathogens induce eIF2α-regulated stress responses in infected, unfed nymphs.** (**A**) Graphic representation of IRE1α-TRAF2 signaling and the Integrated Stress Response pathways in *Ixodes* ticks. (**B-I**) Gene expression in flat, unfed *I. scapularis* nymphs that are either uninfected (-), *A. phagocytophilum*-infected (*A.p.*), or *B. burgdorferi*-infected (*B.b.*). Each data point is representative of 1 nymph. Gene expression was quantified by qRT-PCR using primers listed in Supplemental Table 1. Student’s t-test. *P < 0.05. (**J**) Phosphorylated eIF2α (36 kDa) immunoblot against ISE6 tick cells that were either uninfected (-), infected for 24 hrs (*A. phagocytophilum*: *A.p.*; *B. burgdorferi*: *B.b.;* MOI 50), or treated with the eIF2α phosphorylation inhibitor ISRIB for 1 hr prior to infection (24 hrs). β-actin was probed as an internal loading control (45 kDa). Immunoblots are representative of 2 biological replicates. See also Supplemental Figure 1.

The ISR is a highly conserved signaling network that is activated by cellular stress in eukaryotes^26, 27^. Four different stress-sensing kinases initiate the ISR in mammals: GCN2 (general control nonderepressible), HRI (heme-regulated inhibitor), PKR (protein kinase double-stranded RNA-dependent), and PERK, which is also part of the UPR network^28, 29^. eIF2α is the central regulatory molecule that all ISR kinases converge on, which then activates the transcription factor ATF4 (Fig 1A). ATF4 can also act as a transcriptional repressor of genes that lead to cell death^30, 31^. Although the ISR is much less studied in arthropods relative to mammals, genome analysis demonstrates that ticks encode most ISR components with the exception of a PKR ortholog^18, 21^. We found that *B. burgdorferi* or *A. phagocytophilum* infection transcriptionally induced the ISR kinases (PERK, GCN2, HRI), the eIF2α regulatory molecule, and ATF4 in flat, unfed nymphs (Fig 1E-I).

ISR activation can be monitored by probing for the phosphorylation status of eIF2α^26, 29^. When eIF2α amino acid sequences from human and *I. scapularis* were aligned, we observed a good amount of sequence similarity (Supplemental Figure 1A). Importantly, the activating residue that is phosphorylated by ISR kinases, Ser51, was conserved. We therefore used a commercially raised antibody specific for phosphorylated eIF2α to monitor ISR activation in tick cells. Relative to non-treated controls, ISE6 cells infected with either *A. phagocytophilum* or *B. burgdorferi* showed a band at approximately 36 kDa, correlating with the predicted molecular weight of *I. scapularis* eIF2α (Fig 1J). When tick cells were treated with a small molecular inhibitor of eIF2α phosphorylation, ISRIB (integrated stress response inhibitor)^32^, the 36 kDa band was no longer present, indicating that the band observed was specific to phosphorylated eIF2α (Fig 1J). Altogether, these data show that cellular stress responses converging on eIF2α are activated by *A. phagocytophilum* and *B. burgdorferi* in ticks.

### The PERK pathway promotes A. phagocytophilum growth and survival in tick cells

To determine how eIF2α-regulated stress responses impact pathogen survival in ticks, pharmacological modulators or RNAi silencing were used in *I. scapularis* cells.

ISRIB inhibits phosphorylation of eIF2α^32^, thereby shutting down the ISR. In contrast, salubrinal is an eIF2α activator and promotes ISR activity^33, 34^. We observed that, each pharmacological modulator had opposing effects on *A. phagocytophilum* colonization and replication. Inhibiting eIF2α with ISRIB caused a dose-dependent decline in bacteria (Fig 2A). In contrast, promoting eIF2α activation with salubrinal conferred a survival advantage (Fig 2B). We next used an RNAi-based knockdown approach targeting either *eIF2α* or the downstream transcription factor, *ATF4*. In agreement with pharmacological inhibition, transcriptional silencing caused a decline in *A. phagocytophilum* numbers (Fig 2C-D), indicating that eIF2α-regulated stress responses promote pathogen survival in tick cells.

**Figure 2.**
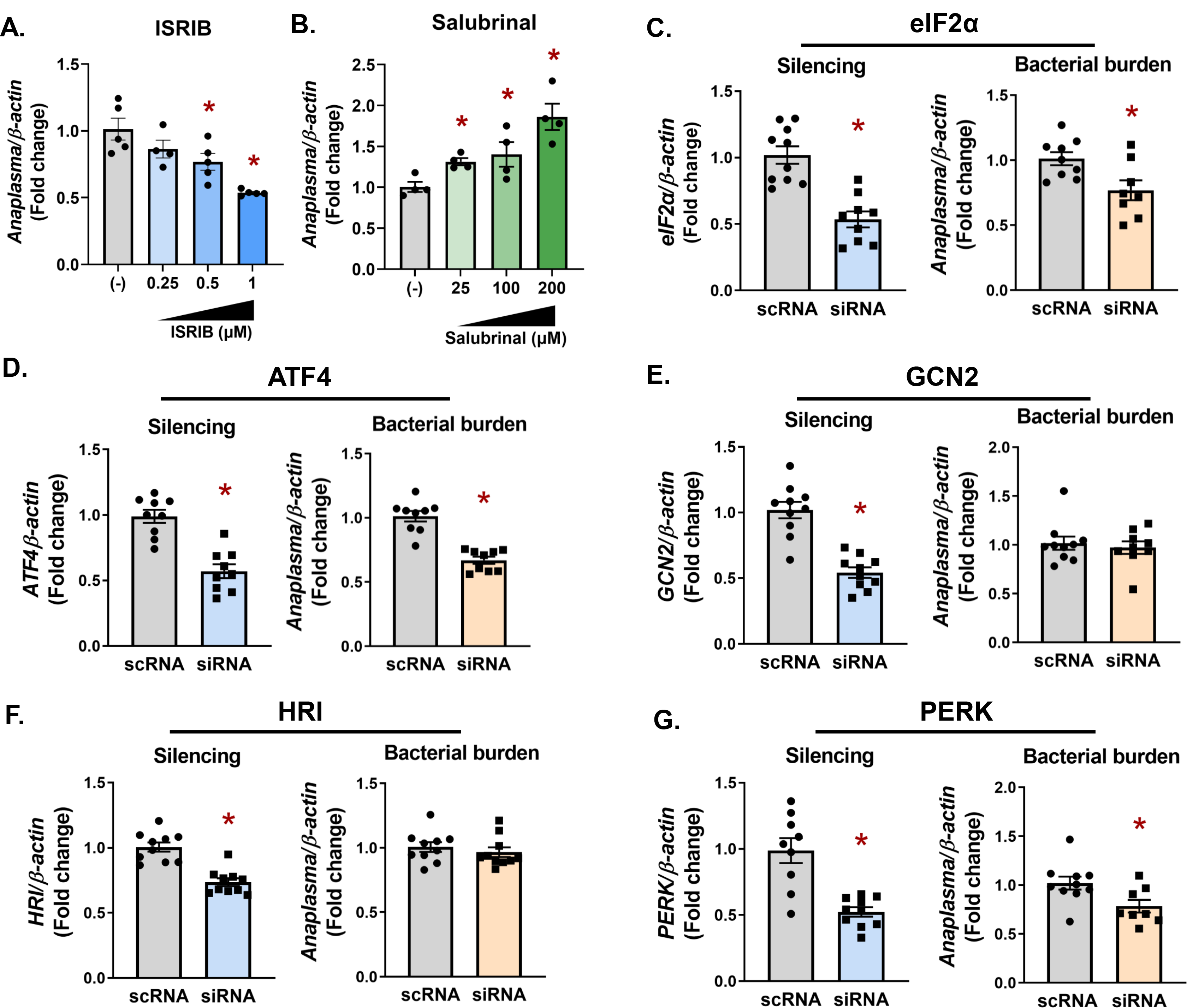
The PERK-eIF2α-ATF4 axis promotes A. phagocytophilum infection in tick cells. (A-B) ISE6 tick cells (1 x 106) were pretreated with ISRIB (A) or salubrinal A. (**B**) at the indicated concentrations for 1 hr prior to infection with *A. phagocytophilum* for 18 hrs (MOI 50). (**C-G**) IDE12 tick cells (1 x 10^6^) were treated with silencing RNAs (siRNA) against indicated genes or scrambled RNA controls (scRNA) for 24 hrs prior to infection with *A. phagocytophilum* (MOI 50) for 18 hrs. *A. phagocytophilum* burden and gene silencing for *eIF2α* (**C**), *ATF4* (**D**), *GCN2* (**E**), *HRI* (**F**), and *PERK* (**G**) were measured by qRT-PCR. Data are representative of at least five biological replicates with at least two technical replicates. Error bars show SEM, *P < 0.05 (Student’s t-test). scRNA, scrambled RNA; siRNA, small interfering RNA.

We next sought to determine which upstream stress-sensing kinase is involved during infection. RNAi knockdown was used to silence the expression of *HRI*, *GCN2*, and *PERK* in tick cells. Although significant silencing was observed for each treatment (Fig 2E-F), a defect in *A. phagocytophilum* survival was only observed with *PERK* knockdown (Fig 2G). This survival defect correlated with what was observed when *eIF2α* and *ATF4* were silenced (Fig 2C-D) or pharmacologically inhibited by ISRIB (Fig 2A), suggesting that PERK may be the activating kinase.

### Pathogen colonization and persistence in ticks is supported by PERK, eIF2α, and ATF4

To determine whether the pro-survival role of the PERK pathway observed *in vitro* had a similar impact on microbes *in vivo,* we used RNAi in *I. scapularis* larvae together with *Anaplasma* or *Borrelia*. Distinct tissue tropisms and kinetics are exhibited in ticks by the intracellular rickettsial bacterium, *A. phagocytophilum*, and the extracellular spirochete, *B. burgdorferi*. *A. phagocytophilum* enters the midgut with a bloodmeal and rapidly traverses the midgut epithelium to colonize the salivary glands^7, 35, 36^. In contrast, *B. burgdorferi* remains in the midgut during the molt and colonizes the tick between the midgut epithelium and peritrophic membrane^37, 38^. Owing to these differences, we evaluated how *Ixodes* PERK signaling impacts colonization and persistence of both pathogens. An overnight siRNA immersion protocol^20^ was used to silence *PERK, eIF2α,* or *ATF4* in *I. scapularis* larvae. The next day, larvae were dried and rested before being placed on infected mice. With this approach, we observed significant knockdown of targeted genes (Fig 3A, E, I; 4A, E, I).

**Figure 3.**
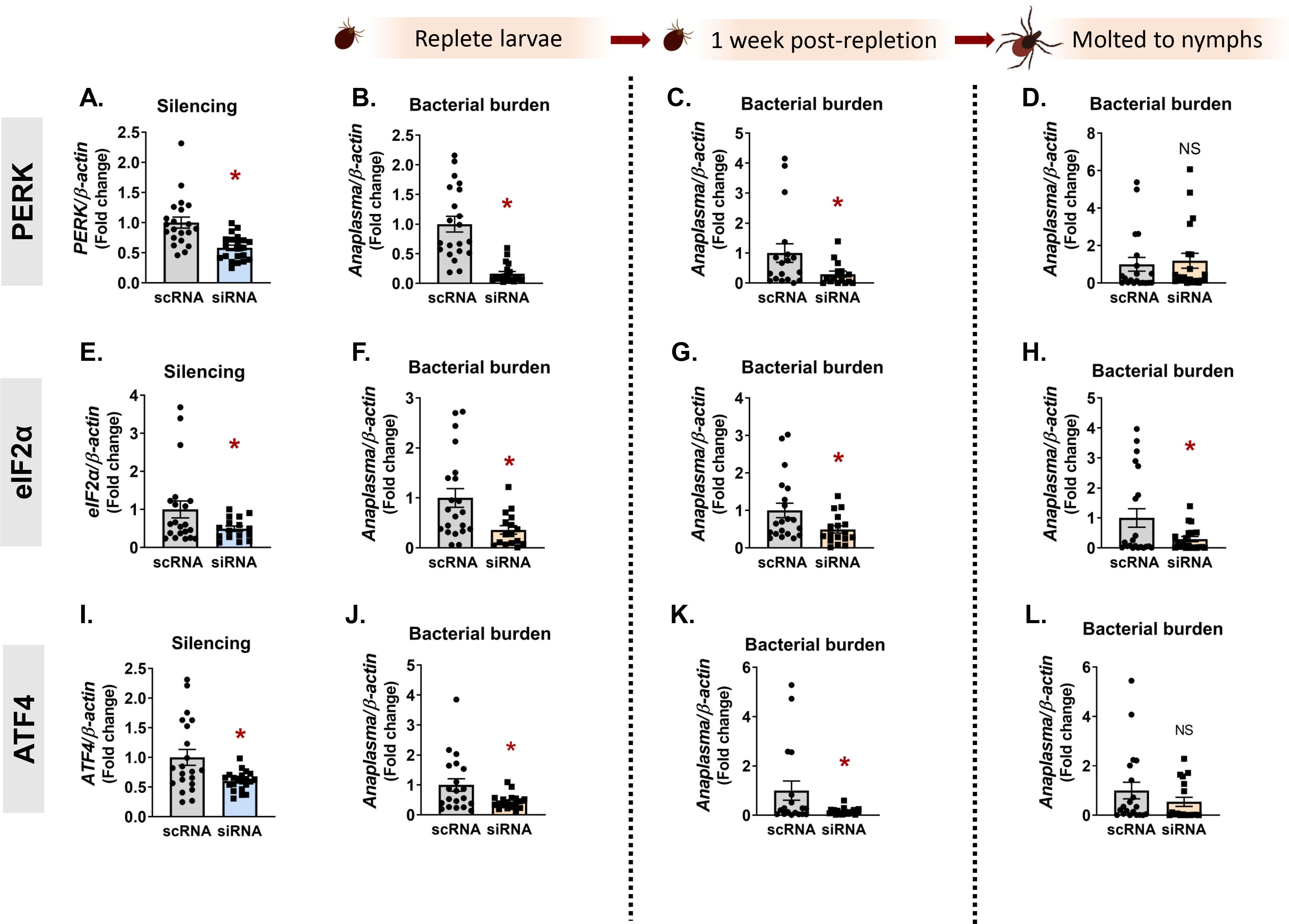
**The PERK pathway supports *A. phagocytophilum in vivo.*** *I. scapularis* larvae were immersed overnight in siRNA targeting *PERK* (**A-D**), *eIF2α* (**E-H**), or *ATF4* (**I-L**) and fed on *A. phagocytophilum*-infected mice. Silencing efficiency (**A, E, I**) and bacterial burden were assessed at three time intervals by qRT-PCR: immediately following repletion (**B, F, J**), one-week post-repletion (**C, G, K**), and after ticks molted to nymphs (**D, H, L**). Data are representative of 10-20 ticks and at least two experimental replicates. Each point represents one tick, with two technical replicates. Error bars show SEM, *p < 0.05 (Welch’s t-test). NS, non-significant. scRNA, scrambled RNA, siRNA, small interfering RNA.

After ticks fed to repletion, pathogen numbers were quantified at three different time points that correspond to: 1) pathogen acquisition (immediately after repletion), 2) population expansion in the tick (7-14 days, post-repletion^39^), and 3) pathogen persistence through the molt (4-6 weeks, post-repletion). Ticks evaluated immediately after repletion (Fig 3B, F, J) and 7 days post-repletion (Fig 3C, G, K) showed a 2-6X reduction in *Anaplasma* numbers, indicating that the PERK-eIF2α-ATF4 pathway has a pro-survival role *in vivo*. However, ticks silenced for *PERK* or *ATF4* as larvae did not show statistically significant differences in *Anaplasma* burden as nymphs (Fig 3D, L).

This may be due to the loss of transcriptional knockdown over the duration of the molt (4-6 weeks) or pathogen numbers rebounding after escaping the midgut to the salivary glands. For *Borrelia*, knocking down *PERK*, *eIF2α*, and *ATF4* (Fig 4A, E, I) also caused a 2-10X decrease in bacterial numbers at early colonization time points (Fig 4B-C, F-G, J-K). However, in contrast to *Anaplasma*, *Borrelia* remained significantly decreased after replete larvae molted to nymphs (Fig 4D, H, L). It is not clear why *Borrelia* remained restricted after the molt while *Anaplasma* did not. One possible explanation is the fundamental difference in tick colonization sites, as the midgut is generally a more hostile environment for invading microbes than the salivary glands^40, 41^. Taken as a whole, these data indicate that the *Ixodes* PERK pathway supports both extracellular and intracellular tick-borne microbes *in vivo*.

**Figure 4.**
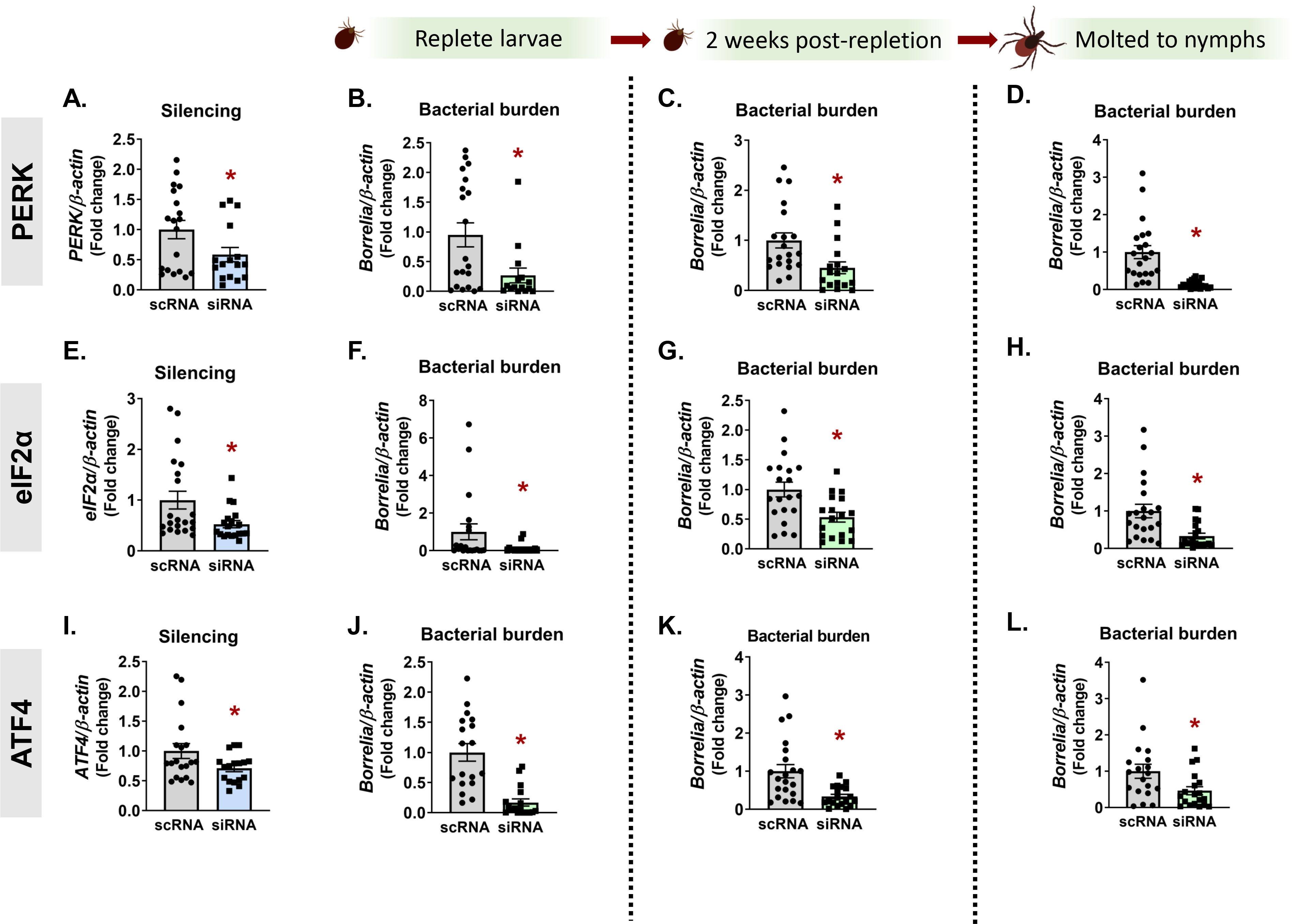
*In vivo B. burgdorferi* colonization and persistence through the molt is supported by the PERK pathway. *PERK* (**A-D**), *eIF2α* (**E-H**), or *ATF4* (**I-L**) were silenced in *I. scapularis* larvae by immersing ticks in siRNA overnight. Recovered ticks were fed on *B. burgdorferi*-infected mice. Silencing efficiency (**A, E, I**) and bacterial burden were assessed at three time intervals by qRT-PCR: immediately following repletion (**B, F, J**), two weeks post-repletion (**C, G, K**), and after ticks molted to nymphs (**D, H, L**). Data are representative of 10-20 ticks and at least two experimental replicates. Each point represents one tick, with two technical replicates. Error bars show SEM, *P < 0.05 (Welch’s t-test). scRNA, scrambled RNA, siRNA, small interfering RNA.

### A. phagocytophilum and B. burgdorferi trigger an Nrf2 antioxidant response in ticks

The microbe-supporting activity of the PERK-eIF2α-ATF4 pathway led us to ask what downstream signaling events occur that functionally promote pathogen survival. Genetic manipulation techniques in *I. scapularis* ticks and tick cell lines remain challenging. To circumvent this limitation, we employed a surrogate reporter system to interrogate downstream signaling events from the PERK pathway. A collection of luciferase reporter plasmids with promoter sequences for transcription factors associated with ER stress (XBP1, NF-κB, CHOP, SREBP1, and Nrf2) were transfected into HEK293 T cells. Transfected cells were then either infected with *A. phagocytophilum* or *B. burgdorferi* or left uninfected. After 24 hrs, luciferase activity was quantified to ascertain the transcriptional activity of each promoter (Fig 5A-B). In agreement with previous reports^20^, XBP1 activation was not observed with either *A. phagocytophilum* or *B. burgdorferi.* In contrast, the immunoregulatory transcription factor NF-κB was significantly induced by both, which is also in agreement with previous findings^42–45^. Infection did not induce CHOP or SREBP1 activity, but did robustly activate the antioxidant regulator Nrf2 (nuclear factor erythroid 2–related factor 2) (Fig 5A-B).

**Figure 5.**
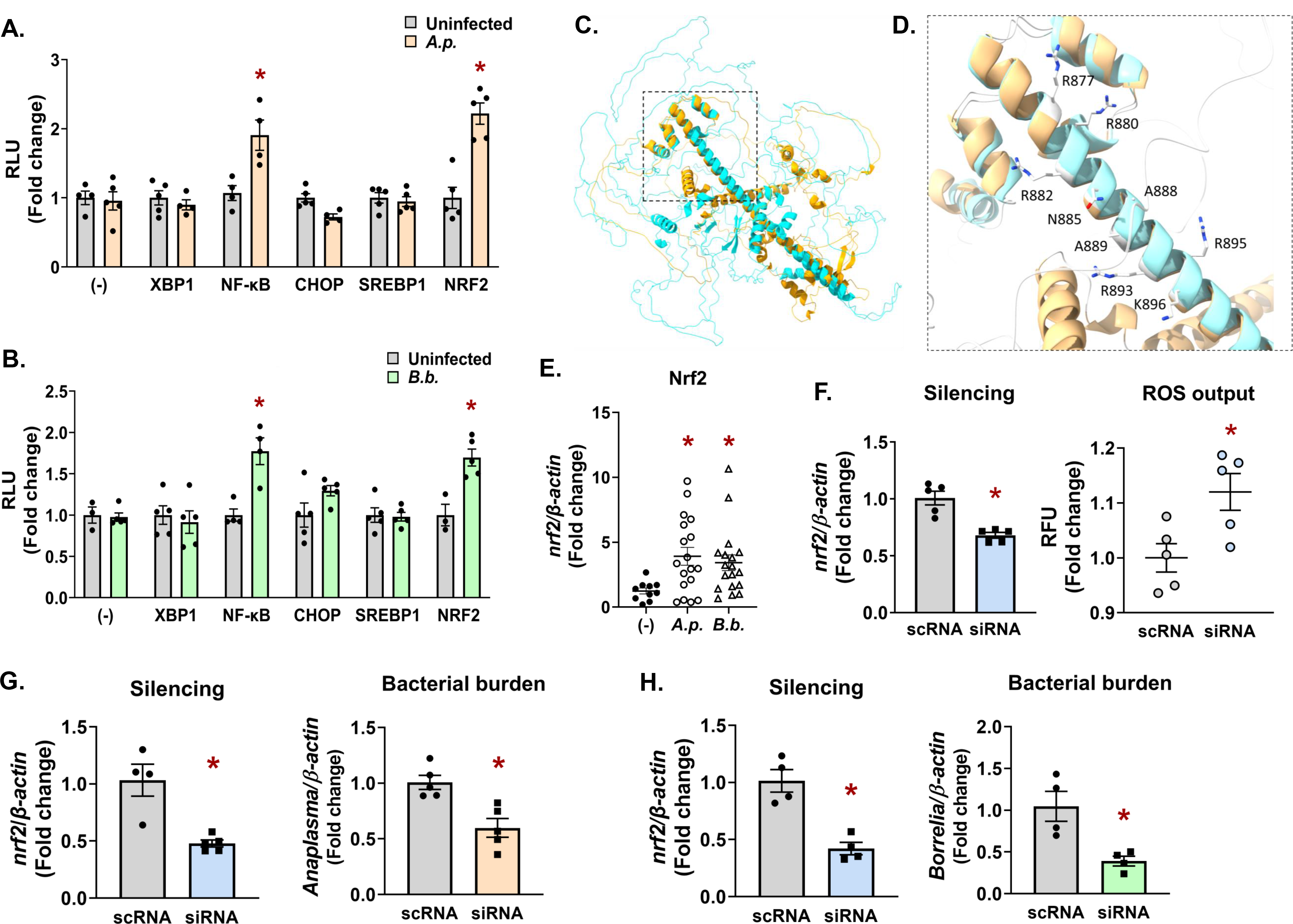
Infection triggers an Nrf2-regulated antioxidant response in ticks that promotes pathogen survival. (**A-B**) HEK293T cells (1 x 10^4^) were transfected with luciferase reporter vectors for assaying activity of ER stress transcription factors XBP1, NF-κB, CHOP, SREBP1, and NRF2 or were untransfected (-). Cells were then infected with *A. phagocytophilum* (*A.p.*) (**A**) or *B. burgdorferi* (*B.b.*) (**B**). After 24 hrs, D-luciferase was added and luminescence was measured as relative luminescence units (RLU). Measurements were normalized to uninfected controls (gray bars). Luciferase assays are representative of 3-5 biological replicates with at least two experimental replicates ± SEM. Student’s t-test. *P < 0.05. (**C-D**) Predicted *Ixodes* Nrf2 structure modeled with AlphaFold^98, 99^ (blue) and overlaid with human Nrf2 (orange) using UCSF ChimeraX^100^. The bZIP domain is indicated by a box with dashed lines. (**D**) Magnified region of the bZIP domain depicting residues that that are predicted to interact with ARE sequences in DNA promoter regions (R877, R880, R882, N885, A888, A889, R893, R895, K896). See also Supplemental Figure 2. (**E**) *Nrf2* expression levels in flat, unfed nymphs that are uninfected (-), *A. phagocytophilum*-infected (*A.p.*), or *B. burgdorferi*-infected (*B.b.*). Each data point is representative of 1 nymph. Gene expression was quantified by qRT- PCR using Nrf2 primers listed in Supplemental Table 1. Student’s t-test. *P < 0.05. (**F- H**) IDE12 tick cells were treated with silencing RNAs (siRNA) targeting *nrf2* for 24 hrs prior to infection with *A. phagocytophilum* (18 hrs) (**F-G**) or *B. burgdorferi* (**H**). Gene silencing (**F-H**) and bacterial burden (**G-H**) were quantified by qRT-PCR. ROS was measured as relative fluorescent units (RFU) after 24 hrs of infection (**F**). Data are representative of at least 4-5 biological replicates and two technical replicates. Error bars show SEM, *P < 0.05 (Student’s t-test). scRNA, scrambled RNA; siRNA, small interfering RNA.

Nrf2 is an evolutionarily conserved cap’n’collar transcription factor that coordinates antioxidant responses^46–50^. It functions by binding to a consensus DNA sequence (antioxidant response elements (ARE)) in the promoter regions of Nrf2- regulated genes^51, 52^. To identify an Nrf2 ortholog in *I. scapularis*, we used the human Nrf2 protein sequence to query the tick genome^53^. A BLAST analysis returned the *Ixodes* protein XP_042149334.1. Although *I. scapularis* Nrf2 had low sequence conservation with human Nrf2 (Supplemental Fig 2A), it did display a high degree of structural conservation when modeled with AlphaFold (Fig 5C-D; *Ixodes* Nrf2- blue; Human Nrf2- orange; Supplemental Fig 2B). Notably, amino acids within the Basic Leucine Zipper (bZIP) domain of Nrf2 that mediate DNA interactions with promoter ARE regions^54^ were well-conserved in the *Ixodes* protein (R877, R880, R882, N885, A888, A889, R893, R895, K896; Supplemental Fig 2A; Fig 5D).

To determine if *nrf2* transcriptionally responds to infection, we evaluated *nrf2* gene expression in flat, unfed nymphs. We observed significantly higher *nrf2* expression in nymphs infected with *A. phagocytophilum* and *B. burgdorferi* relative to uninfected controls (Fig 5E). Since vertebrate Nrf2 regulates basal and inducible antioxidant genes, we next asked whether the *Ixodes* Nrf2 ortholog influences the tick cell redox environment. Tick cells were transfected with silencing RNAs against *nrf2* or with scrambled controls. Cells were then infected with *A. phagocytophilum* and reactive oxygen species (ROS) were measured with the fluorescent indicator 2′,7′- dichlorofluorescein diacetate (DCF-DA). We found that depleting *Ixodes nrf2* caused a significantly higher amount of ROS when compared to scrambled controls (Fig 5F).

ROS is a potent antimicrobial agent and it is well-established that *A. phagocytophilum* and *B. burgdorferi* are sensitive to ROS-mediated killing^55–58^. Considering Nrf2’s role as an antioxidant regulator, we reasoned that silencing *nrf2* expression should enhance microbial killing owing to accumulated ROS. Accordingly, we found that when *nrf2* was knocked down in tick cells, there was a significant decline in *A. phagocytophilum* and *B. burgdorferi* survival (Fig 5G-H). Collectively, these results support the conclusion that *Ixodes* Nrf2 is induced during infection and functionally promotes an antioxidant response, which confers a pro-survival environment for transmissible microbes in the tick.

### Antioxidant activity of the PERK-eIF2α-ATF4 pathway protects pathogen survival in ticks

*A. phagocytophilum* and *B. burgdorferi* induce Nrf2, which is a transcriptional activator downstream from the PERK pathway^27^. We therefore asked whether blocking eIF2α during infection would influence the redox environment in ticks. Tick cells were either uninfected, infected (*A. phagocytophilum* or *B. burgdorferi*), or treated with the eIF2α inhibitor ISRIB prior to infection. Kinetic measurements of ROS and RNS were monitored in tick cells with the fluorescent reporters DCF-DA (ROS) or 4,5- diaminoflurescein diacetate (RNS) (Fig 6A-D). In untreated cells, *A. phagocytophilum* infection caused a rise in ROS that peaked at 24 hrs. Thereafter, ROS levels declined, which is consistent with reports that *A. phagocytophilum* infection suppresses ROS^59–62^. However, when eIF2α signaling is blocked with the ISRIB inhibitor, *A. phagocytophilum* caused increased ROS throughout infection that never declined (Fig 6A; Supplemental Figure 3A). Similarly, *B. burgdorferi* induced ROS in tick cells and treating with ISRIB showed greater accumulation of ROS than infection alone (Fig 6B; Supplemental Figure 3A). Inhibiting eIF2α had similar impacts on RNS in tick cells infected with *A. phagocytophilum* and *B. burgdorferi* (Fig 6C-D; Supplemental Figure 3B). Combining ISRIB with infection conditions caused significantly higher RNS compared to infection alone. Unexpectedly, we also observed that untreated infection conditions showed a decline in RNS, which may suggest that *Anaplasma* and *Borrelia* suppress nitrosative stress in the tick. Collectively, these data indicate that eIF2α signaling functionally coordinates an antioxidant response in tick cells during infection.

**Figure 6.**
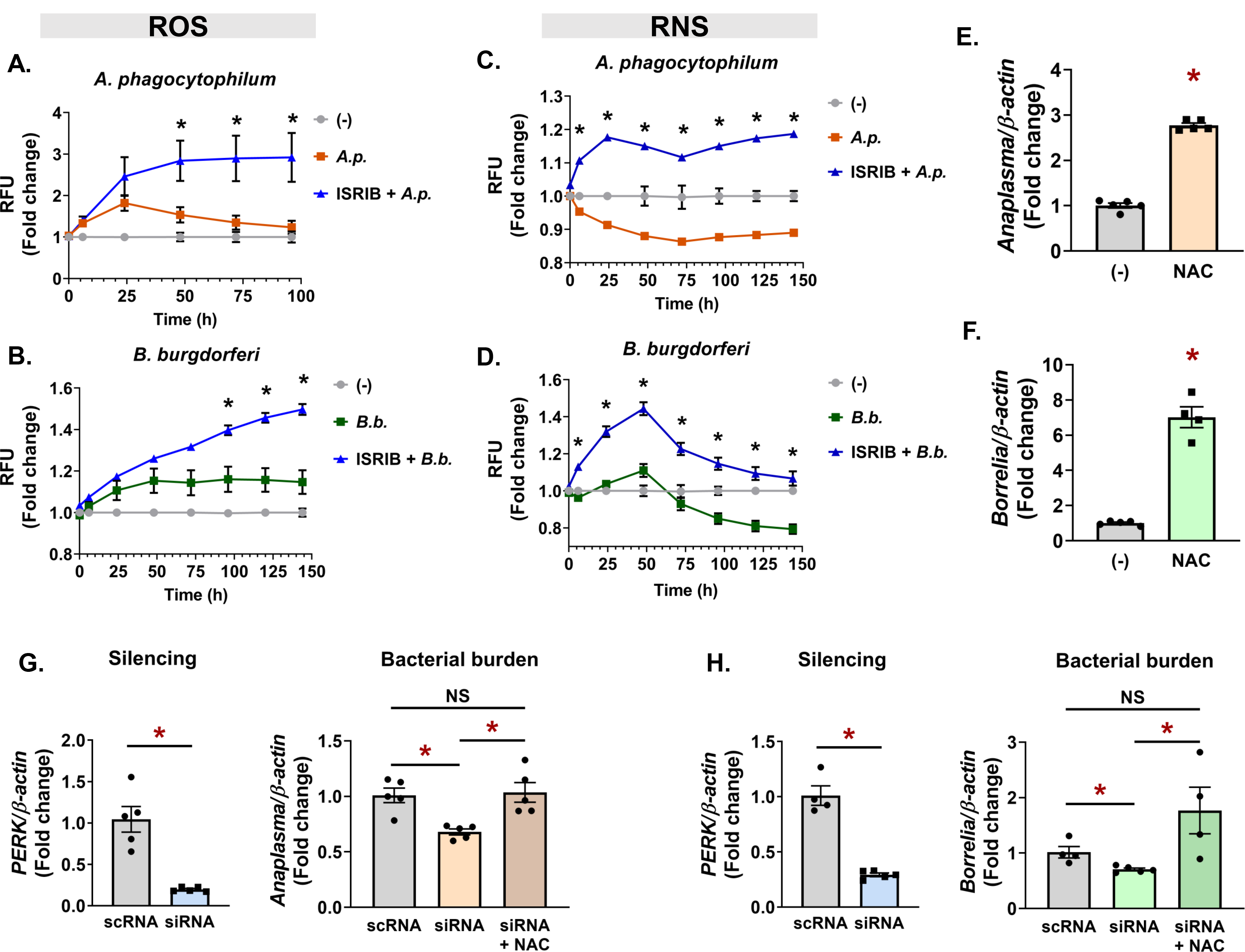
Antioxidant activity of the PERK-eIF2α pathway protects pathogens in ticks. (A-D) ROS (A, B) and RNS (C, D) measurements in ISE6 cells (1.68 x 105) untreated (-), infected (A.p. or B.b.), or pretreated with 1µM ISRIB prior to infection with A. *phagocytophilum* (ISRIB + *A.p.*) (**A, C**) or *B. burgdorferi* (ISRIB + *B.b.*) (**B, D**). Fluorescence was measured at the indicated time points and is presented as RFU, normalized to untreated, uninfected controls (-). (**E-F**) IDE12 cells were infected with *A. phagocytophilum* (**E**) or *B. burgdorferi* (**F**) alone or in the presence of 5mM NAC for 24 hrs. (**G-H**) *perk* was silenced in IDE12 cells (1 x 10^6^). Cells were infected with *A. phagocytophilum* (**G**) or *B. burgdorferi* (**H**) alone or in the presence of 5mM NAC. Silencing levels and bacterial burdens were quantified by qRT-PCR. Data are representative of at least 4-5 biological replicates and two technical replicates. Error bars show SEM, *P < 0.05 (Student’s t-test). NAC, N-acetyl cysteine. scRNA, scrambled RNA; siRNA, small interfering RNA.

We next asked if the antioxidant environment potentiated by the PERK-eIF2α- ATF4 pathway was the functional mechanism that supports pathogen survival in ticks. We first established that antioxidants enhance microbial survival in ticks. Tick cells that were supplemented with the antioxidant N-acetyl cysteine (NAC) during infection showed significantly more *A. phagocytophilum* or *B. burgdorferi* survival when compared to untreated controls (Fig 6E-F). We then asked if the antioxidant activity of NAC could rescue the microbicidal phenotype caused by silencing *perk*. Tick cells were treated with silencing RNA against *perk* or scrambled controls, then infected with *A. phagocytophilum* or *B. burgdorferi* with and without NAC. As previously observed (Fig 2G), silencing the expression of *perk* caused a significant decrease in pathogen survival. However, supplementing with exogenous antioxidants rescued the bactericidal effect caused by blocking the PERK pathway (Fig 6G-H). Altogether, our findings support a model where transmissible pathogens activate the PERK-eIF2α-ATF4 pathway, which functionally supports pathogen persistence in ticks through an Nrf2- mediated antioxidant response (Fig 7).

**Figure 7.**
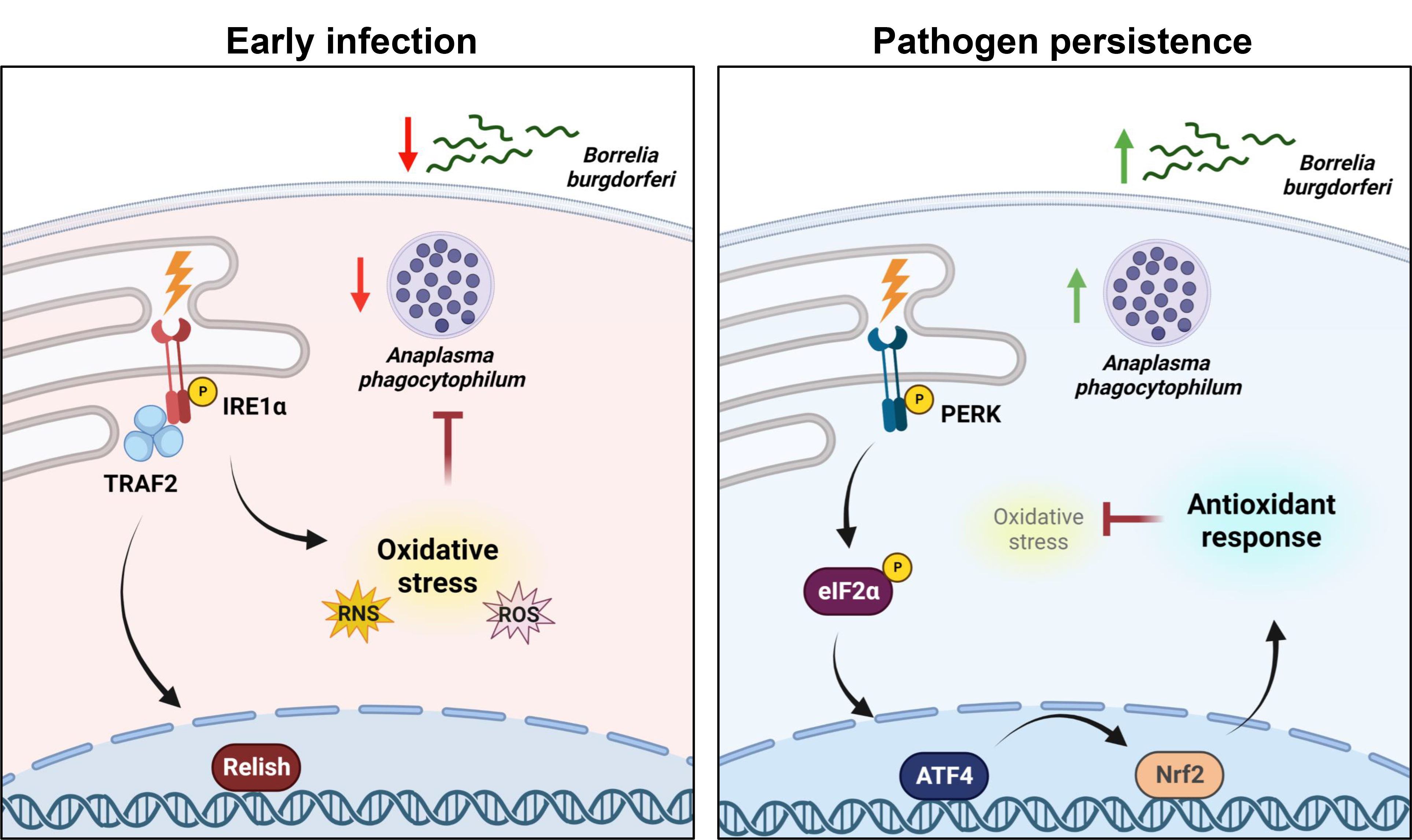
The PERK-eIF2α-ATF4 axis promotes pathogen survival in ticks through an Nrf2-mediated antioxidant response. When colonizing the tick, *A. phagocytophilum* and *B. burgdorferi* trigger the *Ixodes* IMD pathway and ROS/RNS through the IRE1α-TRAF2 axis of the UPR. Tick-borne microbes persist in the tick over time by stimulating the PERK branch of the UPR, which signals through eIF2α and the transcription factors ATF4 and Nrf2 to trigger an antioxidant response that promotes microbial survival.

## DISCUSSION

How pathogens persist in the tick is likely a multifaceted topic involving complex interactions orchestrated by both the microbe and the arthropod. In this article, we shed light on one aspect of this subject by demonstrating that *Anaplasma* and *Borrelia* infection activates the *Ixodes* PERK-eIF2α-ATF4 stress response pathway, which facilitates pathogen survival. The microbe-benefiting potential of this pathway was ultimately connected to an antioxidant response that is mediated by the *Ixodes* Nrf2 ortholog. Collectively, our findings have uncovered a piece of the puzzle in understanding how pathogens can persist in the tick despite immunological pressure from the arthropod vector.

*A. phagocytophilum* and *B. burgdorferi* have significantly different lifestyles (obligate intracellular vs. extracellular) and tissue tropisms (salivary glands vs. midgut), but both induce a state of oxidative stress upon tick colonization^20^. Given that these microbes are susceptible to killing by oxidative and nitrosative stress^55–58, 62–67^, it is perhaps not surprising that both would benefit from an antioxidant response in the tick. However, some discrepancy between *Anaplasma* and *Borrelia* survival phenotypes was observed at different time points *in vivo.* Bacterial colonization was decreased in larvae when *PERK*, *eIF2α*, or *ATF4* were knocked down by RNAi (Fig 3-4). In contrast, *Borrelia* remained significantly reduced in molted nymphs, but *Anaplasma* numbers rebounded. This may be attributable to differences in tissue tropisms for each pathogen. *Anaplasma* rapidly escapes the midgut and colonizes the salivary glands^7, 35, 36^, whereas *B. burgdorferi* remains in the midgut during the molt^37, 38^. The midgut is a niche that is generally hostile to microbes owing to several factors including ROS and RNS production^40, 41, 64, 65^ and may explain why *Borrelia* numbers are restricted even after larvae molt to nymphs. The *Ixodes* salivary gland environment also produces ROS and RNS^64^, but antioxidant proteins found in salivary glands, such as the periredoxin Salp25D^68^, may protect *Anaplasma* and explain why pathogen numbers rebounded after the molt.

Since *A. phagocytophilum* and *B. burgdorferi* are susceptible to oxidative and nitrosative damage^55–58, 62–67^, these microbes may be inducing the *Ixodes* PERK-eIF2α- ATF4-Nrf2 pathway to create a more hospitable environment and facilitate persistence. *A. phagocytophilum* replicates intracellularly and secretes a suite of effectors that manipulate host cell biology and promote the formation of a replicative niche. Although only one tick-specific effector has been characterized to date^69^, it is conceivable that *Anaplasma* manipulates PERK pathway activation in the tick with secreted effector molecules. *B. burgdorferi* replicates extracellularly and does not encode any secretion systems for effector transport, which makes direct host-cell manipulation less-likely. However, it is possible that *Borrelia* may transport small molecules^63^ that could activate the PERK pathway and promote an antioxidant response. Alternatively, the PERK- eIF2α-ATF4-Nrf2 signaling cascade may be responding to general stress signals caused by infection^70^. For example, pathogens can secrete toxic by-products, compete with the host for limiting amounts of nutrients, and/or cause physical damage to host cells^25^. Our previous study demonstrated that both *Borrelia* and *Anaplasma* activate the IRE1α-TRAF2 branch of the UPR in ticks, which results in the accumulation of ROS^20^. When this pathway was inhibited, ROS levels were either partially (*Anaplasma*) or completely (*Borrelia*) mitigated. Since oxidative stress is an important stimuli that triggers the UPR^71, 72^, it is possible that ROS potentiated by the IRE1α-TRAF2 pathway is the signal that activates the PERK pathway at later time points and results in an antioxidant response. From this perspective, the PERK-eIF2α-ATF4-Nrf2 pathway may be a host-driven response that promotes the preservation of “self”.

Unexpectedly, we observed that *A. phagocytophilum* and *B. burgdorferi* caused a decline in RNS that began either a few hours after infection (*A.p.)* or after 2 days (*B.b.*) (Fig 6C-D). A potential explanation for this could be increased Arginase expression.

Arginase competes for the nitric oxide synthase substrate L-arginine and is therefore a potent inhibitor of nitric oxide production^73^. Villar *et al* reported that *Ixodes arginase* expression levels are significantly increased in *A. phagocytophilum-*infected ticks^74^, which could explain the rapid decline in RNS we observed after *Anaplasma* infection (Fig 6C). Similarly, a recent report by Sapiro *et al* analyzed *I. scapularis* nymphs by mass spectrometry and reported that Arginase was enriched with *B. burgdorferi* after 4 days of feeding, but not at early time points^75^. This may explain why RNS also decreased with *B. burgdorferi* (Fig 6D), but only after 48 hours of infection.

The Nrf2 gene regulatory network has not yet been characterized in *I. scapularis*. Mammalian Nrf2 regulates components of the glutathione and thioredoxin antioxidant systems as well as enzymes involved in NADPH regeneration^76, 77^. Given that ROS levels increased in tick cells when Nrf2 was knocked down (Fig 5F), it is reasonable to speculate that similar antioxidant genes are regulated by *Ixodes* Nrf2. Moreover, it is well-established that tick-borne microbes benefit from antioxidant gene expression in the tick^68, 78–85^. For example, manipulating selenium-related antioxidant gene expression has microbicidal consequences for microbes in the tick^78–80, 83, 85, 86^. This is in agreement with our findings that Nrf2 expression promotes *Borrelia* and *Anaplasma* survival (Fig 5G-H) and further indicates that the *Ixodes* Nrf2 coordinates an antioxidant gene network.

Between human and *Ixodes* Nrf2, we observed structural conservation in the bZIP domain with 100% conservation of the amino acids that make direct contact with DNA (Fig 5C-D). However, there was very low sequence conservation (Supplemental Fig 2A) and the *Ixodes* Nrf2 is almost 400 amino acids longer than human Nrf2. This may suggest that there are regulatory mechanisms or protein-protein interactions that are unique to the tick. In addition to protein differences, the *Ixodes* Nrf2-regulated gene network also appears to be divergent. For example, heme oxygenase is an important cytoprotective protein regulated by Nrf2 in most eukaryotes and has been implicated in disease tolerance^87^. However, chelicerates do not have a gene encoding heme oxygenase^88^. Altogether, this suggests that there are differences in Nrf2 and the genes it coordinates between ticks and other eukaryotes, which may be tailored to the life histories of each organism. The extent of divergence between *Ixodes* Nrf2 and other eukaryotes is a question that remains unanswered at this time.

Collectively, our findings illustrate a scenario where early tick infection triggers IREα-TRAF2 signaling leading to IMD pathway activation and ROS production^20^, while persistent infection induces the PERK pathway and an antioxidant response through Nrf2 that supports pathogen survival (Fig 7). Innate immune mediators, such as AMPs and ROS, have potent antimicrobial activity but the non-specificity of these molecules can also cause damage to host tissues^70^. We speculate that the PERK-driven antioxidant response in persistently infected ticks is a host-driven response aimed at reducing collateral damage to “self”. Ultimately this network preserves tick fitness, but also promotes pathogen persistence. The result is a balance between microbial restriction and host preservation that promotes arthropod tolerance to infection by transmissible pathogens.

## METHODS

### Bacteria and animal models

Roswell Park Memorial Institute (RPMI) 1640 medium supplemented with 10% heat-inactivated fetal bovine serum (FBS; Atlanta Biologicals, S11550) and 1x Glutamax (Gibco, 35050061) was used to culture *A. phagocytophilum* strain HZ in HL60 cells (ATTC, CCL-240). Cultures were maintained between 1 x 10^5^-1 x 10^6^ cells/ml at 37°C in the presence of 5% CO2. Mice were infected with 1 x 10^7^ host cell free *A. phagocytophilum* in 100µl of PBS (Intermountain Life Sciences, BSS-PBS) intraperitoneally as previously described^20, 89^. Six days post-infection 25-50µl of infected blood was collected from the lateral saphenous vein of each mouse and *A. phagocytophilum* burdens assessed via quantitative PCR (*16s* relative to mouse β- actin^20, 90, 91^).

*B. burgdorferi* B31 (MSK5^20, 92^) was grown at 37°C with 5% CO2 in modified Barbour-Stoenner-Kelly II (BSK-II) medium supplemented with 6% normal rabbit serum (NRS; Pel-Freez, 31126-5). Density and growth phase of the spirochetes were assessed by dark-field microscopy. Prior to infection, plasmid verification was performed as previously described^20, 92^. Mice were inoculated with 1 x 10^5^ low passage spirochetes in 100µl of 1:1 PBS:NRS intradermally. Mice were bled from the lateral saphenous vein at 7d post-infection. 25-50µl of *B. burgdorferi*-infected blood was cultured in BSK-II medium and examined for the presence of spirochetes by dark-field microscopy^20, 93, 94^.

Male C57BL/6 mice, aged 6-10 weeks old, obtained from colonies maintained at Washington State University were used for all experiments. Guidelines and protocols approved by the American Association for Accreditation of Laboratory Animal Care (AAALAC) and by the Office of Campus Veterinarian at Washington State University (Animal Welfare Assurance A3485-01) were used for all experiments utilizing mice. The animals were housed and maintained in an AAALAC-accredited facility at Washington State University in Pullman, WA. All procedures were approved by the Washington State University Biosafety and Animal Care and Use Committees.

*Ixodes scapularis* ticks at the larval stage were obtained from the Biodefense and Emerging Infectious Diseases (BEI) Research Resources Repository from the National Institute of Allergy and Infectious Diseases (www.beiresources.org) at the National Institutes of Health or from Oklahoma State University (OSU; Stillwater, OK, USA).

Ticks were maintained in a 23°C incubator with 16:8h light:dark photoperiods and 95- 100% relative humidity.

### Tick cell and HEK293 T cultures

*I. scapularis* embryonic cell lines ISE6 and IDE12 were cultured at 32°C with 1% CO2 in L15C-300 and L15C media, respectively. These growth media were supplemented with 10% heat-inactivated FBS (Sigma, F0926), 10% tryptose phosphate broth (TBP; BD, B260300), and 0.1% lipoprotein bovine cholesterol (LPBC; MP Biomedicals, 219147680)^20, 95^.

HEK293 T cells were maintained in Dulbecco’s modified Eagle medium (DMEM; Sigma, D6429) supplemented with 10% heat-inactivated FBS (Atlanta Biologicals; S11550) and 1x Glutamax. Cells were maintained in T75 culture flasks (Corning; 353136) at 33°C or 37°C in 5% CO2.

### Pharmacological treatments and RNAi silencing

ISE6 and IDE12 cells were seeded at 1 x 10^6^ cells per well in a 24-well plate and pretreated with ISRIB (Cayman Chemical, 16258), or salubrinal (Thermo Scientific, AAJ64192LB0) for 1h prior to infection. Cells were infected with *A. phagocytophilum* or

*I. B. burgdorferi* at an MOI of 50 for 18h alone or in the presence of 50mM N-acetyl cysteine (NAC; Sigma, A7250). Cells were collected in RIPA buffer (for immunoblotting) or TRIzol for RNA (Invitrogen, 15596026). RNA was extracted with the Direct-zol RNA Microprep Kit (Zymo; R2062) and cDNA was synthesized from 300-500ng total RNA using the Verso cDNA Synthesis Kit (Thermo Fisher Scientific, AB1453B). Bacterial burden was assessed by qRT-PCR with iTaq universal SYBR Green Supermix (Bio- Rad, 1725125). Cycling conditions used were as recommended by the manufacturer.

Transfection experiments used siRNAs and scrambled controls (scRNAs) synthesized with the Silencer siRNA Construction Kit (Invitrogen, AM1620). ISE6 or IDE12 cells were seeded 1 x 10^6^ cells per well in a 24-well plate or 2.5 x 10^5^ per well in a 96-well plate. 3µg of siRNA or scRNA in conjunction with 2.5µl Lipofectamine 2000 (Invitrogen, 11668027) were transfected into tick cells overnight in 24-well plates. 1µg of siRNA or scRNA with 1µl Lipofectamine 2000 was used for 96-well plates. Cells were infected with *A. phagocytophilum* (MOI 50) or *B. burgdorferi* (MOI 50) for 18h. Cells infected with *Anaplasma* had the cell culture supernatant removed before collecting in TRIzol. Cells infected with *Borrelia* had both cells and supernatant collected in TRIzol. RNA was isolated and transcripts assessed by qRT-PCR as described above. All data are expressed as means ± SEM.

### Polyacrylamide gel electrophoresis and immunoblotting

Protein concentrations from cells collected in RIPA buffer were quantified by bicinchoninic acid (BCA) assay (Pierce; 23225). 25µg of protein was loaded onto a 4- 15% MP TGX precast cassette (Bio-Rad; 4561083) and proteins were separated at 100V for 1 h 25 min. Proteins were transferred to a polyvinylidene difluoride (PVDF) membrane and were blocked with 5% BSA (bovine serum albumin) in TBS-T (1x tris-buffered saline containing 0.1% Tween 20) for 1 to 2 h at room temperature. The eIF2α antibody (1:500; EMD Millipore 07-760-I) was incubated with the PVDF membrane overnight at 4°C in 5% BSA in TBS-T. The following day a secondary antibody was applied (donkey anti-rabbit–HRP; Thermo Fisher Scientific; A16023; 1:2,000). Blots were visualized with enhanced chemiluminescence (ECL) Western blotting substrate (Thermo Fisher Scientific; 32106).

### ROS and RNS assays

1.68 x 10^5^ ISE6 cells per well were seeded in a 96-well plate with black walls and clear bottoms (Thermo Scientific, 165305). All wells were treated with the fluorescent detection probes 2’,7’-dichlorofluorescein diacetate (10µM, DCF-DA; Sigma, D2926) or 4,5-diaminoflurescein diacetate (5µM, DAF-2DA; Cayman Chemical, 85165) for 1h in Ringer Buffer (155mM NaCl, 5mM KCl, 1mM MgCl2 · 6H2O, 2mM NaH2PO4 · H2O, 10mM HEPES, and 10mM glucose) ^20, 96^. Cells were treated with the probe alone or in the presence of 1µM ISRIB. Buffer was removed and cells washed with room temperature PBS. *A. phagocytophilum* or *B. burgdorferi* were then added at an MOI of 50 in the presence of ISRIB or vehicle control (DMSO). Fluorescence was measured at 504nm/529nm at the indicated times and data are graphed as fold change of relative fluorescence units (RFU) normalized to the negative control ± SEM.

### Luciferase reporter assay

HEK293 T cells were seeded in white-walled, clear-bottom 96-well plates (Greiner Bio-One, 655098) at a density of 1 x 10^4^ cells per well. The following day, cells were transfected with 0.05µg of each vector from the UPR/ER stress response luciferase reporter vector set (Signosis, LR-3007) and 0.5µl of Lipofectamine 2000 in Opti-MEM I reduced-serum medium (Gibco, 31985062). Transfections were allowed to proceed overnight. The following day, the medium containing the plasmid-Lipofectamine 2000 complex was removed and replaced with complete DMEM for an additional 18- 24h. Cells were then infected with *A. phagocytophilum* MOI 50 or *B. burgdorferi* at an MOI of 200 or left uninfected overnight. The following day, D-luciferin potassium salt (RPI, L37060) was added to each well at a final concentration of 5mg/ml and luminescence measured. Data are graphed as relative light units (RLU) normalized to uninfected controls ± SEM.

### Gene expression analysis of whole ticks

Gene expression profiling of whole ticks was performed on flat, unfed nymphs that were infected with *A. phagocytophilum* or *B. burgdorferi* as larvae. Individual ticks were snap frozen in liquid nitrogen and mechanically pulverized prior to the addition of TRIzol. RNA extraction and qRT-PCR analysis was performed as described above with primers listed in Supplemental Table 1. Gene expression levels were measured by qRT-PCR and normalized to uninfected controls. Data are expressed as means ± SEM.

### RNAi silencing and analysis of whole ticks

RNAi silencing in *I. scapularis* larvae was performed as described previously^20^. Briefly, approximately 150 larvae were transferred to a 1.5ml tube with 40µl of siRNA or scrambled controls and incubated overnight at 15°C. Larvae were then dried and allowed to recover overnight under normal maintenance conditions prior to being placed onto mice the following day. Larvae were allowed to feed to repletion and frozen at three time points: immediately following collection, after resting (7d for *A. phagocytophilum*, 14d for *B. burgdorferi*), and after molting into nymphs. Replete larvae were weighed in groups of three to assess feeding efficiency before being processed individually, as described above. qRT-PCR analysis was performed with the use of a standard curve to generate absolute numbers of the target sequences. Primers used to generate the plasmids used in the standard curves are the same as the primers used to measure target levels (Supplemental Table 1), with the exception of *A. phagocytophilum 16S*.

### Protein alignments and modeling

*I. scapularis* proteins were identified using NCBI (National Center for Biotechnology Information) protein BLAST and querying the tick genome with human protein sequences for PERK (NP_004827.4) and Nrf2 (NP_001138884.1). Alignments were visualized with JalView^97^. Physiochemical property conservation between amino acids is indicated by shading. AlphaFold^98, 99^ was used to model the protein structure of *Ixodes* Nrf2 and align it to the human Nrf2 protein structure. Alignments were visualized with UCSF ChimeraX^100^.

### Statistical analysis

*In vitro* experiments were performed with 3-5 replicates. *In vivo* experiments used at least 10-20 ticks. Data were expressed as means ± SEM and analyzed with either an unpaired Student’s t-test or Welch’s t-test. Calculations and graphs were created with GraphPad Prism. A P-value of < 0.05 was considered statistically significant.

## ACKNOWLEDGEMENTS

We are grateful to Ulrike Munderloh (University of Minnesota) for providing ISE6 and IDE12 tick cell lines; Jon Skare (Texas A&M Health Science Center) for providing *B. burgdorferi* B31 (MSK5); BEI Resources and Oklahoma State University for *Ixodes scapularis* ticks, and Arden Baylink (Washington State University) for guidance with AlphaFold and UCSF ChimeraX. Schematics in Figures 1 and 7 were created with Biorender.com.

## Author contributions

K.L.R., J.H., and D.K.S. designed the study. K.L.R., J.H., E.A.F., K.A.V., A.L.W., L.C.S- L., S.J.W., E.R-Z., J.M.P., and D.K.S. contributed to methodology, investigation, and data analysis. All authors provided intellectual input into the study. K.L.R. and D.K.S. wrote the manuscript; all authors contributed to editing.

