## Supplemental Figure 1 for "PERK-mediated antioxidant response is key for pathogen persistence in ticks"

**Supplemental Figure 3 | ISRIB does not potentiate ROS or RNS in tick cells.**

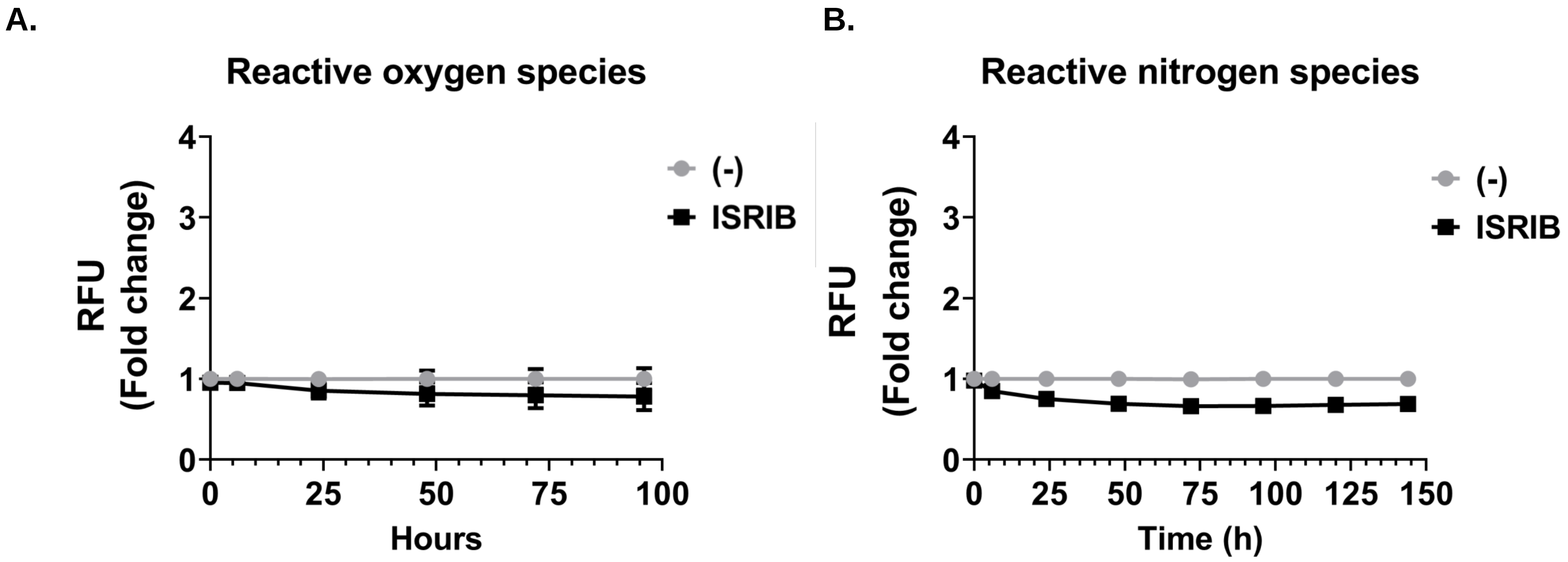

**Supplemental Figure 3 | ISRIB does not potentiate ROS or RNS in tick cells.** ROS (**A**) and RNS (**B**) measurements in ISE6 cells (1.68 x $10^5$). Cells were untreated (-) or pretreated with 1µM ISRIB. Fluorescence was measured at the indicated time points and is presented as RFU, normalized to untreated controls (-).
