## Supplemental Figure 2 for "PERK-mediated antioxidant response is key for pathogen persistence in ticks"

**Supplemental Figure 1 | Sequence alignment of eIF2α from *I. scapularis* and *Homo sapiens*.**

## A.

```
H.s._eIF2α    1 MPGLSCRFYQHKFPEVEDVVMVNVRSIAEMGAYVSLLEYNNIEGMILLSELSRRRIRSINKLIRIGRNECVVVIRVDKEKGYIDLSKRRVSPEEAIKCEDKFTKSKTVYS 110
I.s._eIF2α    1 -MPISCRFYSQKYPEPDDVVMVNVRQIAEMGAYVRLLEYNDIEGMILLSELSRRRIRSINKLIRVGRNECVVVIRVDKDKGYIDLSKRRVSPEDITKCEEKFARGKAVNS 109

H.s._eIF2α  111 ILRHVAEVLEYTKDEQLESLFQRTAWVFDDKYKRPGYGAYDAFKHAVSDPSILDSLDLNEDEREVLINNINRRLTPQAVKIRADIEVACYGYEGIDAVKEALRAGLNCST 220
I.s._eIF2α  110 ILRHVAETLKYETDAQLEELYKKTAWYFDEKFKKT-TAAYDVFKLAVTDLSVLDECDLDPNTREVLLTNIKHRLTPQAVKIRADIEVSCYEYEGIDAVKHALREGMALST 218

H.s._eIF2α  221 ENMPIKINLIAPPRYVMTTTTLERTEGLSVLSQAMAVIKEKIEEKRGVFNVQMEPKVVTDTDETELARQMERLERENAEVDGDDDAEEMEAKAED------------- 315
I.s._eIF2α  219 EDMPIKINLIAPPVYVMTTTTMDRTEGLDALNAAIVKIEETIKKAGGVFTVQNPPKVVTETDELELAKQLDRLEQANAEIDGDDDDEEAEAGSGSEAGEDDNVVIAEA 326
```

**Supplemental Figure 1 | Sequence alignment of eIF2α from *I. scapularis* and *Homo sapiens*.** Available protein sequences from NCBI were imported into Jalview to visualize sequence alignments. Shaded amino acid residues indicate conservation index and percentage identity between the two proteins. The activity-inducing phosphoserine of eIF2α, Ser51, is indicated with black shading.
