## Supplemental Figure 3 for "PERK-mediated antioxidant response is key for pathogen persistence in ticks"

**Supplemental Figure 2 | *Ixodes* Nrf2 sequence alignment and structural prediction with AlphaFold.**

**A.**

```
H.s._Nrf2        -------------------------------------------------------------------------------------------------------------------
I.s._Nrf2      1 MMNYKKCSGVRSTILQLLLAVSLLQLGSNPDSNPVRVDRRSDLVRSVRMGVHSAFTQTQSLSPPVFVHPKNVDAFDADPIFAELHELGIPSLLRRIRTRVFAYLAEDPGLPAVKA  115

H.s._Nrf2      1 ----------------------MMDL---ELPPPGLPSQQDMDLIDILWRQDIDLGVSREVFDFSQRRKEYELEKQ----------KKLEKERQEQLQKEQEKAFFAQLQLDEET   80
I.s._Nrf2    116 SGDAPTPANTGQQQRATSGDPVIDDASSESTGSGADLTQEDMDLIEILWKQDEDLGVPRDVFDFRFEGNDSGDATIKEQLESTEVPTKMENDSLSELPDDSNPWEGFQYSIDSET  230

H.s._Nrf2     81 GEFL--------------------PIQPAQHIQS-ETSGSANYSQVAH-----IPKSDALYF-------------DDCM-------QLLAQ-------TFPFVDDNEVSSAT---  139
I.s._Nrf2    231 GEHILGDADKRGASRERTDSGCCLPMDEDTGLSLPEEEQEALFAELDRILQGQSPQAADLYPVEENWTCPEPRPGDDACGRGLQCGTLGAGGPPPHNYTDPFPPGVLLHNATLGP  345

H.s._Nrf2    140 --FQSLVPDIPGHIE------------------------SPVFIATNQAQSPETSVAQV--------------APVDLD--------------------GMQQDIEQVWEELLSI  194
I.s._Nrf2    346 PPADHMAPFMHGSFGTTACTGLPCNDGSSLGEGPPYGQDFPDYLYPPTSG-MGGTVGTVGGLNMGMCVGNLSGPPVGAHGQNDYLYTDFLGDEDLHLMDMVASPSEGEWTGLLPF  459

H.s._Nrf2    195 PELQCLNIENDKLVETTMVPSPEAKLTEVDN-YHFYSSIPSME----------KEVGNCSPH-FLNAF-----------------------EDSFSSILSTEDPNQLTVNSLNSD  274
I.s._Nrf2    460 GVLVWSGST----AGPKKRPQPAADLYPVEENWTCPEPRPGDDACGRGLQCGTLGAGGPPPHNYTDPFPPGVLLHNATLGPPPADHMAPFMHGSFGTTACTGLPCN-DGSSLGEG  569

H.s._Nrf2    275 ATVNTDFGDE----------------------------------------FYSAFIAEPSISNSMPSPATLSHSLSELLNGPIDVSDLSLCKAFNQNHPESTAEFNDSDSGISLN  349
I.s._Nrf2    570 PPYGQDFPDYLYPPTSGMGGTVGTVGGLNMGMCVGNLSGPPVGAHGQNDYLYTDFLGDEDLHLM-----DMVASPSEAASGPYPVQ-LSVAE---------DRTDNSSDTGVSSM  669

H.s._Nrf2    350 TS---PSVASPEHSVES----S----SYGD---TLLGLSDSEVEELDSAP-GSVKQ------NG-----PK------TPVHS-SGDM------------------VQPLSPSQGQ  413
I.s._Nrf2    670 GSERVPSISEDHEWLDSCSESSSHADGEGGDLPRFLGSAQKKYKLFGRKPPGPGENDTVVLGVGGGYPCPAPPHHYPVPVHRLPGGATVALVPKPASAPNFPSPHIPRLAPPVGA  784

H.s._Nrf2    414 S-THVHDAQCENTPEKELPVSPGHR--------KTPFTKDKHSSRLEAHLTRDELRAKALHIPFPVEKIINLPVVDFNEMMSKEQFNEAQLALIRDIRRRGKNKVAAQNCRKRKL  519
I.s._Nrf2    785 LVQHNHSYSAAEAA--SSTAEEGNTSDEDEEDGGCELGEAERLRQDQLRASRDEKRARALNLPLTNVEIVELPIEEFNERLAKFELTEAQLALIRDIRRRGKNKVAAQNCRKRKL  897

H.s._Nrf2    520 ENIVELEQDLDHLKDEKEKLLKEKGENDKSLHLLKKQLSTLYLEVFSMLRDEDGKPYSPSEYSLQQTRDGNVFLVPKS------------KKPDVKKN-----------       605
I.s._Nrf2    898 DQIMSLQQDVEALHLQRQELERRHEELLA-QRLLG---RDKYSRLCQLLATHATRPLSPGSEQVAAL--GTPLLDAATLANGASDDAQALRGARLKKKAAVDIKWEEDE       1000
```

**B.**

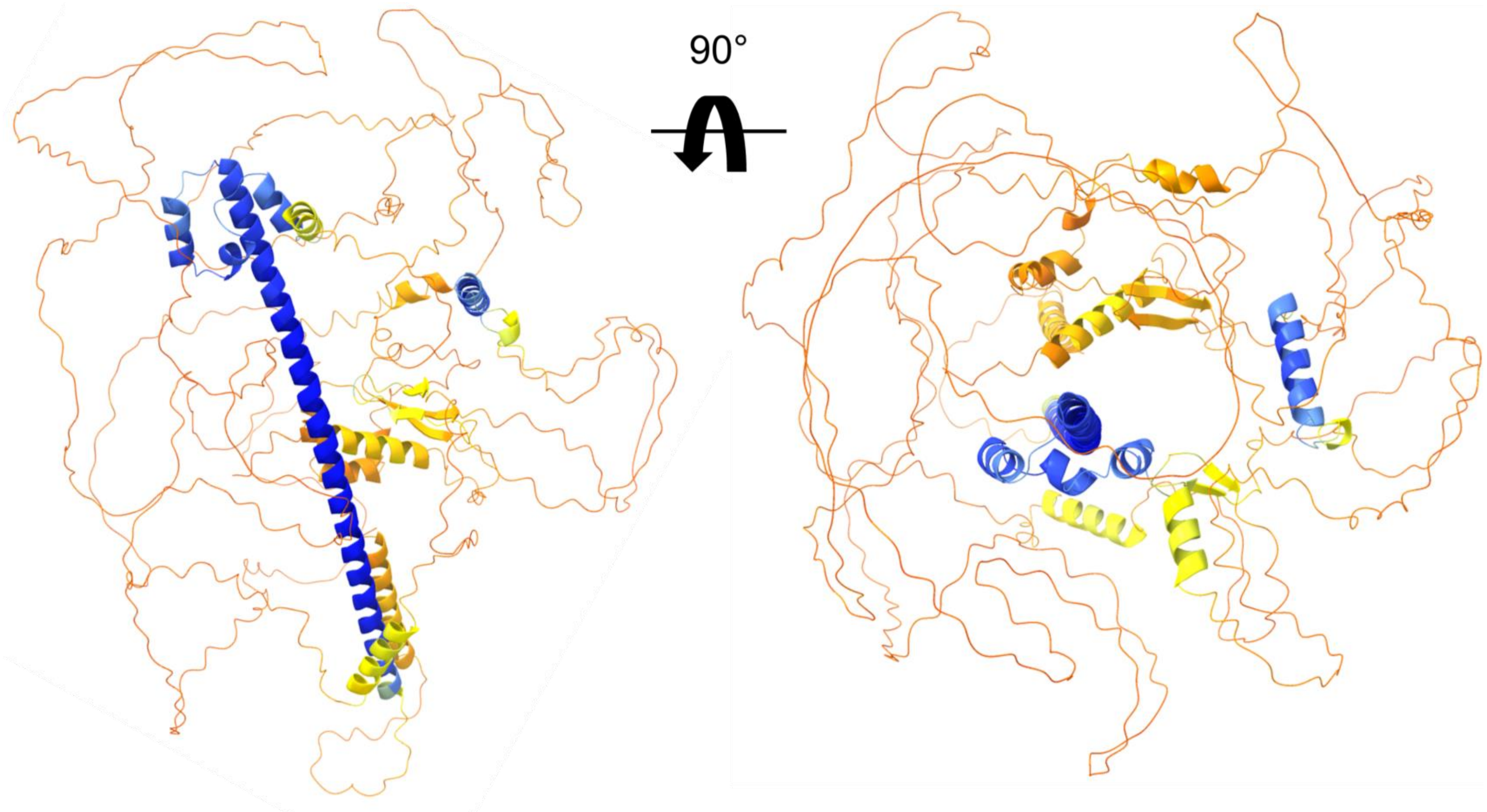

**Supplemental Figure 2 | *Ixodes* Nrf2 sequence alignment and structural prediction with AlphaFold.** (**A**) Human and *Ixodes* Nrf2 protein sequences were aligned and visualized with Jalview. The conservation index and percentage identity between the two proteins is indicated by shaded amino acid residues. Amino acids that mediate DNA interactions with promoter ARE regions are indicated with black shading. (**B**) *Ixodes* Nrf2 protein as predicted by AlphaFold. Each residue is color-coded based on the model confidence score, pLDDT. Blue indicates the most confidently predicted regions. Orange to yellow indicates regions of low confidence.
