## Supplemental Table 1 for "PERK-mediated antioxidant response is key for pathogen persistence in ticks"

| Name | Target gene | Primer Sequences |
|---|---|---|
| *Mus musculus* β-Actin (qRT-PCR) | XM_030254057.1 | F 5'-ACGCAGAGGGAAATCGTGCGTGAC-3'<br>R 5'-ACGCGGGAGGAAGAGGATGCGGCAGTG-3' |
| *Anaplasma phagocytophilum* 16S (qRT-PCR) | NC_007797 | F 5'-CCCTAAGGCCTTCCTCACTC-3'<br>R 5'-CAGCCACACTGGAACTGAGA-3' |
| *Anaplasma phagocytophilum* 16S_full | NC_007797 | F 5'- TCCTGGCTCAGAACGAACG-3'<br>R 5'- GTCACTGACCCAACCTTAAATGG-3' |
| *Borrelia burgdorferi* FlaB (qRT-PCR) | MN954474.1 | F 5'-TTGCTGATCAAGCTCAATATAACCA-3'<br>R 5'-TTGAGACCCTGAAAGTGATGC-3' |
| *Ixodes scapularis* Actin (qRT-PCR) | XM_029977298.1 | F 5'-GCCGGGACCTTACAGACTATC-3'<br>R 5'-CACGGACAATTTCACGCTCG-3' |
| *Ixodes scapularis* IRE1α (qRT-PCR) | XM_029972190.1 | F 5'-GAGAAGGCCATCTTCGTCGG-3'<br>R 5'-GAGTAGCCTGGGCAGCATAG-3' |
| *Ixodes scapularis* TRAF2 (qRT-PCR) | XM_029977983.1 | F 5'-CCGCGAAAAGAACAGCTTAC-3'<br>R 5'-TACCACGTTGGACTCCTTCC-3' |
| *Ixodes scapularis* BiP (qRT-PCR) | XM_002433611.2 | F 5'-ATCGTGTGGTGTCTAGCGG-3'<br>R 5'-CGATACCGATGACTGTGCCG-3' |
| *Ixodes scapularis* PERK (qRT-PCR) | XM_029994352.4 | F 5'-ATCCACTCCTGTACTTGGGC-3'<br>R 5'-TTCACCCGATTCTGAAGGCT-3' |
| *Ixodes scapularis* HRI (qRT-PCR) | XM_029992945.4 | F 5'-TATTCCGAAGCTGTCTCCGC-3'<br>R 5'-GTCAGCAGGTTGTAGCCCTA-3' |
| *Ixodes scapularis* GCN2 (qRT-PCR) | XM_029969993.4 | F 5'-GCCCCAAGAAAGCATGACTC-3'<br>R 5'-TGCTTCGTTGGGATACTGGT-3' |
| *Ixodes scapularis* eIF2α (qRT-PCR) | XM_040212362.2 | F 5'-ATCCGGTCCATCAACAAGCT-3'<br>R 5'-TCCTTGTCCACCCTGATGAC-3' |
| *Ixodes scapularis* ATF4 (qRT-PCR) | XM_029967345.4 | F 5'- AGGTTGTCTACTGCCCGTAC-3'<br>R 5'- TCCCAGTCGACTTCCATGTC-3' |
| *Borrelia burgdorferi* cp9 (PCR) | BBC10 | F 5'-GAACTATTTATAATAAAAAGGAGAGC-3'<br>R 5'-ATCTTCTTCAAGATATTTTATTATAC-3' |
| *Borrelia burgdorferi* cp26 (PCR) | BBB19 | F 5'-AATAATTCAGGGAAAGATGGG-3'<br>R 5'-AGGTTTTTTTGGACTTTCTGCC-3' |
| *Borrelia burgdorferi* lp17 (PCR) | BBD10 | F 5'-CAAACTTATCAAATAGCTTATC-3'<br>R 5'-ACTGCCACCAAGTAATTTAAC-3' |
| *Borrelia burgdorferi* lp25 (PCR) | BBE16 | F 5'-ATGGGTAAAATATTATTTTTTGGG-3'<br>R 5'-AAGATTGTATTTTGGCAAAAAATTTTC-3' |
| *Borrelia burgdorferi* lp28-1 (PCR) | BBF20 | F 5'-ATGAACAAAAAAATTTTCTATTTC-3'<br>R 5'-GTTGCTTTTGCAATATGAATAGG-3' |
| *Borrelia burgdorferi* lp28-2 (PCR) | BBG02 | F 5'-TCCCTAGTTCTAGTATCTACTAGACCG-3'<br>R 5'-TTTTTTTTGTATGCCAATTGTATAATG-3' |
| *Borrelia burgdorferi* lp28-3 (PCR) | BBH06 | F 5'-GATGTTAGTAGATTAAATCAG-3'<br>R 5'-TAATAAAGTTTGCTTAATAGC-3' |
| *Borrelia burgdorferi* lp28-4 (PCR) | BBI16 | F 5'-CAGGCCGGATTTTAATATCGA-3'<br>R 5'-GTTTATATTTTGACACTATAAG-3' |
| *Borrelia burgdorferi* lp36 (PCR) | BBK19 | F 5'-AAGTTTATGTTTATTATTGC-3'<br>R 5'-ATTGTTAGGTTTTTCTTTTCC-3' |
| *Borrelia burgdorferi* lp38 (PCR) | BBJ34 | F 5'-AAATTCTATGGAAGTGATG-3'<br>R 5'-TTTCTATTTATTTTTAGGC-3' |

| | | |
|---|---|---|
| *Borrelia burgdorferi* Ip54 (PCR) | BBA16 | F 5'-GCACAAAAAGGTGCTGAG-3'<br>R 5'-TTTTAAAGCGTTTTTAAGC-3' |
| *Borrelia burgdorferi* Ip56 (PCR) | BBQ56 | F 5'-AAGATTGATGCAACTGGTAAAG-3'<br>R 5'-CTGACTGTAACTGATGTATCC-3' |
| *Ixodes scapularis* eIF2α _siRNA_438 | XM_040212362.2 | F 5'-AACCGGAAGACATCACGAAATCCTGTCTC-3'<br>R 5'-AAATTTCGTGATGTCTTCCGGCCTGTCTC-3' |
| *Ixodes scapularis* eIF2α_ scRNA | N/A | F 5'-AAGCATAGCGGAACCTAACAACCTGTCTC-3'<br>R 5'-AATTGTTAGGTTCCGCTATGCCCTGTCTC-3' |
| *Ixodes scapularis* ATF4_siRNA_863 | XM_029967345.4 | F 5'-AAGCAGAGTCCTTTCCGGAAACCTGTCTC-3'<br>R 5'-AATTTCCGGAAAGGACTCTGCCCTGTCTC-3' |
| *Ixodes scapularis* ATF4_ scRNA | N/A | F 5'-AAGGCACAGCTTCGCAGTAATCCTGTCTC-3'<br>R 5'-AAATTACTGCGAAGCTGTGCCCCTGTCTC-3' |
| *Ixodes scapularis* PERK_siRNA_1064 | XM_029994352.4 | F 5'-AAGCATAGAATGGAAGCCCTACCTGTCTC-3'<br>R 5'-AATAGGGCTTCCATTCTATGCCCTGTCTC-3' |
| *Ixodes scapularis* PERK_scRNA | N/A | F 5'-AAGGTAACAAACCGGAGTCATCCTGTCTC-3'<br>R 5'-AAATGACTCCGGTTTGTTACCCCTGTCTC-3' |
| *Ixodes scapularis* HRI_siRNA_231 | XM_029992945.4 | F 5'-AAGGATCATACTCCTGATCTTCCTGTCTC-3'<br>R 5'-AAAAGATCAGGAGTATGATCCCCTGTCTC-3' |
| *Ixodes scapularis* HRI_scRNA | N/A | F 5'-AAACTATTCGTCTACTCGGTACCTGTCTC-3'<br>R 5'-AATACCGAGTAGACGAATAGTCCTGTCTC-3' |
| *Ixodes scapularis* GCN2_siRNA_372 | XM_029969993.4 | F 5'-AACCAACACAATACACCTCAACCTGTCTC-3'<br>R 5'-AATTGAGGTGTATTGTGTTGGCCTGTCTC-3' |
| *Ixodes scapularis* GCN2_scRNA | N/A | F 5'-AAACCAACATACCATAACACCCCTGTCTC-3'<br>R 5'-AAGGTGTTATGGTATGTTGGTCCTGTCTC-3' |
| *Ixodes scapularis* Nrf2_siRNA_1058 | XM_042293400.1 | F 5'-AACCTTCATGCATGGATCCTTCCTGTCTC-3'<br>R 5'-AAAAGGATCCATGCATGAAGGCCTGTCTC-3' |
| *Ixodes scapularis* Nrf2_scRNA | N/A | F 5'-AAGCTTACCGAGTCTCCTATTCCTGTCTC-3'<br>R 5'-AAAATAGGAGACTCGGTAAGCCCTGTCTC-3' |
| *Ixodes scapularis* Nrf2 (qRT-PCR) | XM_042293400.1 | F 5'-GTCTTCGACTTCCGGTTTGA-3'<br>R 5'-GTAGGCACTTCGGTGCTCTC-3' |
